# A One Health study of *Klebsiella pneumoniae* species complex plasmids shows a highly diverse and ecologically adaptable plasmidome

**DOI:** 10.1101/2025.09.14.676155

**Authors:** Mia A Winkler, Marit A K Hetland, Håkon Pedersen Kaspersen, Ragna-Johanne Bakksjø, Eva Bernhoff, Aasmund Fostervold, Jane Hawkey, Bjørn-Tore Lunestad, Nachiket P Marathe, Niclas Raffelsberger, Ørjan Samuelsen, Marianne Sunde, Arnfinn Sundsfjord, Margaret M C Lam, Iren H Löhr

**Author notes:** Corresponding author: Mia A. Winkler. These authors contributed equally (Margaret M C Lam, Iren H Löhr).

## Abstract

Plasmids play a pivotal role in the horizontal gene transfer of antimicrobial resistance (AMR) and virulence determinants among bacteria. Members of the *Klebsiella pneumoniae* species complex (KpSC) can colonise humans, animals, and various environments, and frequently cause nosocomial and community-acquired infections in humans. While plasmid-borne AMR genes are prevalent in clinical strains, the diversity, distribution, and association of plasmids encoding AMR and virulence across ecological niches remain poorly characterised. Understanding the traits governing successful plasmid transmission within and between ecological niches is critical for developing effective AMR prevention strategies. Here, we characterise the diversity and distribution of KpSC plasmids and identify potential niche associations of AMR, heavy metal resistance, virulence factors, and plasmid clusters.

We analysed the plasmidome (i.e. total genetic content attributable to plasmids) of 578 whole-genome sequenced KpSC isolates collected in Norway between 2001-2020 from human (n=453), terrestrial animal (n=102), and marine (n=23) sources. Plasmids from complete hybrid assemblies were annotated and clustered to evaluate the plasmid diversity and content across niches. Additionally, the representativeness of this plasmid collection was determined by clustering with a global collection of 8656 circularised KpSC plasmids.

In total, 1415 circularised plasmids were identified and grouped according to rearrangement distance using Pling, resulting in 130 clusters (containing >1 plasmid), of which 36% (n=47) contained plasmids from >1 niche. The plasmids exhibited significant diversity, as 37% (n=524) remained singletons after clustering. AMR and virulence genes existed across diverse clusters and singletons, but predominantly resided on 120-250 kbp conjugative or mobilisable plasmids harbouring various transposable elements.

The human niche exhibited a significantly higher prevalence of plasmids with AMR genes compared to animal or marine niches (p<0.001), whereas the animal niche displayed a significantly higher prevalence of virulence-encoding plasmids compared to human or marine niches (p<0.001), which was largely due to an enrichment of *iuc*3 plasmids in pigs.

The high diversity of the KpSC plasmids underscores the dynamic nature of plasmid evolution, driven by horizontal gene transfer, and selective pressures. The presence of variable clusters, marked by high genetic diversity, indicates a dynamic plasmidome capable of rapid adaptation to environmental pressures through the acquisition and rearrangement of accessory genes.

## Introduction

*Klebsiella pneumoniae* is a frequent cause of nosocomial and community-acquired infections (1). Its ability to proliferate in humans, animals, and various terrestrial and marine environments underscores its adaptability and the pervasive challenge it poses to public health systems (1–4). The *K. pneumoniae* species complex (KpSC) encompasses seven closely related subspecies that can be further divided into sublineages (SLs) and sequence types (STs) based on genetic similarity (5–7). Clones associated with multidrug resistance (MDR) frequently harbour acquired antimicrobial resistance (AMR) genes that encode extended-spectrum β-lactamases (ESBLs) and carbapenemases, the majority of which are plasmid-borne and spread through horizontal gene transfer (HGT) (1,8,9). Hypervirulence-associated (HV) clones typically harbour genes encoding siderophores, aerobactin and salmochelin, and hypermucoidity co-residing on distinct plasmids (1,8,10).

Plasmids serve as the primary vectors facilitating the transfer of AMR, virulence, and heavy metal resistance (HMR) genes amongst bacteria via HGT (11–14). KpSC isolates exhibit a diverse assortment of plasmids, and often harbour multiple plasmids per isolate. The flexibility of the KpSC accessory genome reflects its promiscuity and capacity to acquire plasmids from a wide variety of species, making KpSC members efficient conduits for HGT. This adaptability positions KpSC as a reservoir for plasmids, facilitating recombination events amongst plasmids which may further promote the spread of clinically relevant genes (1,11). Consequently, understanding the KpSC plasmidome is necessary for the development of strategies aimed at curtailing the spread of AMR and virulence. Despite the central role of plasmids and other mobile genetic elements (MGEs), the factors enabling their successful spread and persistence in KpSC across SLs and ecological niches remain poorly understood. Recent advances in long-read sequencing technologies and hybrid assemblies now enable detailed analyses of complete plasmid sequences, including structural variation and single nucleotide polymorphisms (SNPs) (15–17).

Here, we have utilised a comprehensive collection of 1415 closed plasmids from 578 hybrid-assembled genomes (15) to examine the diversity of KpSC plasmids across ecological niches. The genomes were representative of a larger collection of 3255 Norwegian KpSC isolates, sampled over two decades (2001-2020) from human, terrestrial animal (simply “animal” hereafter), and marine sources (4,18–26). To our knowledge, this is one of the largest scale analyses of the KpSC plasmidome using long-read sequencing within a One Health framework. Through cross-sectorial genotyping and plasmid clustering, we examined plasmid diversity across SLs and ecological niches and identified AMR and virulence determinants within plasmid clusters.

## Materials and Methods

### Sample selection and whole-genome sequencing

The isolates included in this study were selected as a subset of a larger collection of 3255 KpSC isolates collected in Norway between 2001 and 2020 from human infection (21,26) and colonisation (20) (n=2655, from blood, urine, and faeces), animals (19,24,25) (n=500, from broilers, dogs, pigs, and turkeys), and marine sources (18,22,23) (n=99, from bivalves and seawater). All isolates were short-read sequenced on Illumina MiSeq or HiSeq 2500 platforms, and 578 isolates (17.8% of 3255) were selected for additional long-read sequencing on Oxford Nanopore Technologies (ONT) platforms (4,27). Isolates were selected based on KpSC diversity (i.e. to represent the diversity in the pangenome and at least one of each niche-overlapping SL) and the presence of clinically relevant genetic features, including AMR and virulence genes and plasmid replicon markers (see Supplementary Methods for details). Hybrid assembly was performed using both long-read-first and short-read-first approaches on all sequences to produce closed genomes and maximise plasmid recovery. Sequencing and assembly methods are described in detail in Hetland et al. (27).

### Pangenome estimation and genotyping

All genomes were annotated with Bakta v1.8.1 (28) with database v5.0 using the “complete” flag (otherwise default settings) and homologous genes were clustered using Panaroo v1.3.3 (29). The Bakta output was used to identify HMR and thermoresistance genes (see Supplementary Methods). Kleborate v2.4.0 (30) was used to identify species and ST, as well as virulence and AMR determinants on each replicon. MOB-typer v3.1.5 (31) was used to assign primary MOB cluster IDs as well as predict plasmid mobility, GC content, MOB type, and replicon markers. Transposable elements (TEs) were detected using Mobile Element Finder v1.1.2 (32). SLs were defined previously (4) using the *Klebsiella* BIGSdb-Pasteur web tool (https://bigsdb.pasteur.fr/klebsiella/) to assign life identification number (LIN) code prefixes based on core genome multilocus sequence type (cgMLST) profiles (7).

The copy number of each plasmid contig (complete and incomplete, n=1427) was estimated by mapping raw short and long reads to their corresponding closed genomes using Plassembler v1.8.0 (33) in “assembled” mode using the ‘--skip_mash’ flag and ‘-- depth_filter’ set to 0.05, otherwise default settings (database download 2025-07-10). Estimated copy number was assigned as follows: the value of ‘plasmid_copy_number_long’ was used for 1) plasmids ≥20 kpb and 2) plasmids <20 kpb and prepared with the ONT rapid kit; ‘plasmid_copy_number_short’ was used for plasmids with a mean long read depth of zero and plasmids <20 kbp prepared with the ONT ligation kit. Values less than one were rounded up to one and values greater than one were rounded to the nearest decimal. The plasmid load of an isolate was defined as the percentage of total genomic content attributable to plasmids, calculated by multiplying the length of each plasmid by its estimated copy number and dividing the total plasmid nucleotide content by the copy number-adjusted genome size. All plasmid contigs were used to determine the number of distinct plasmids per isolate and the total plasmid load of each isolate.

### Plasmid clustering

#### Norwegian plasmids

The circularised plasmid sequences (n=1415) from our collection were clustered with Pling v2.0 (17) using a batch size of 750 and default settings for other parameters. Briefly, plasmids within the containment distance threshold (i.e. the proportion of shared sequence between two plasmids relative to the smaller of the two, default=0.5) were grouped into communities. Plasmids were then further split into subcommunities (hereafter “clusters’) based on the maximum number of structural rearrangements separating two plasmid sequences (default=4), determined by the Double Cut and Join insertion-deletion (DCJ-indel) distance. Hub plasmids were defined as plasmids that were highly connected to other sparsely or non-connected plasmids or plasmid groups (default: node degree > 10, neighbouring node’s edge density <0.2), which may erroneously connect unrelated plasmids resulting in overclustering (17) (see Supplementary Methods for more details). In addition to the hub plasmids detected by Pling, clusters were manually inspected and additional plasmids were deemed hubs if they appeared to spuriously connect otherwise disparate plasmid groups within a cluster based on characteristics such as sequence length, AMR or virulence gene content, replicon markers, or MOB classification. All identified hub plasmids were excluded from the final network visualisation.

#### Global plasmids

To assess whether our collection of Norwegian plasmids were representative of the global distribution of KpSC plasmids, we retrieved publicly available complete genomes from NCBI’s RefSeq database on 2024-12-11. The search date range was from 2015-01-01 to 2024-12-11 and included all KpSC members. The 578 genomes analysed in this study were removed from the search to avoid duplicate sequences. All genomes were analysed using Kleborate v2.4.0 to confirm species assignment and identify AMR and/or virulence genes. Publicly available genomes were then excluded if: 1) the chromosome was not complete (circularised); 2) the species reported in the RefSeq database was not reported or was not in agreement with the species reported by Kleborate; 3) Kleborate reported a “weak” species match; and/or 4) sample geographical location was not reported in the RefSeq database. Of the remaining genomes, only complete, circularised plasmid sequences were extracted (n=8656) and combined with the plasmids from our dataset, resulting in n=10072 plasmids from 65 countries in six world regions. These were clustered with Pling using the same parameters as described above, to assess the representativeness of our Norwegian KpSC plasmid collection.

## Statistical analysis

Statistical analyses were performed in R version 4.3.1 (2023-06-16); comparisons were made using Kruskal-Wallis (overall) and Mann-Whitney (pairwise) tests for range and chi-squared tests for proportions (overall and pairwise). P-values <0.05 were considered statistically significant and reported as follows: ^*^ P<0.05, ^**^ P<0.01, ^***^P<0.001, ^****^P<0.0001, ns P≥0.05.

## Definitions

Multidrug resistant (MDR): plasmids with a Kleborate num_resistance_classes value ≥3; defined by presence of AMR genes and mutations identified by Kleborate.

## Results

A total of 568/578 (98.3%) whole-genome sequenced KpSC isolates in this study were fully closed (i.e. circularised chromosomes and plasmids). The 578 genomes represented the human (78.4%; n=453/578), animal (17.6%, n=102), or marine niche (4.0%, n=23), and the majority belonged to *K. pneumoniae sensu stricto*, hereafter *K. pneumoniae* (85.1%, n=492), followed by *K. variicola* subsp. *variicola*, hereafter *K. variicola* (11.9%, n=69), *K. quasipneumoniae* subsp. *similipneumoniae* (1.6%, n=9), *K. quasipneumoniae* subsp. *quasipneumoniae* (1.2%, n=7), and a single *K. quasivariicola* isolate (0.2%). The 578 genomes belonged to 291 SLs, the majority of which were represented by a single isolate (72.2%, n=210). The most prevalent SLs in this subset were SL17 (n=24, 4.2%), SL3010 and SL37 (each n=23, 4.0%), SL107 (n=21, 3.6%), SL35 (n=16, 2.8%), SL45 (n=15, 2.6%), and SL258 (n=11, 1.9%), which largely reflected the SL prevalence in the broader collection (4).

Of the 578 genomes, 88.9% (n=514) contained plasmids. In total, 1427 plasmids were present in the genome collection: 1415 circularised, one linear as confirmed by comparing with a set of previously defined linear plasmids (34), and 11 were not fully closed. All plasmid sequences (n=1427) were used to assess plasmid burden across niches, while the 1415 circularised plasmids were used in plasmid clustering and characterisation.

### Plasmid load varied by niche

The distribution of plasmids closely reflected the number of isolates from each niche: the majority (75.8%, n=1082/1427) were present in isolates from the human niche, followed by the animal (18.1%, n=258/1427) and marine (6.1%, n=87/1427) niches (Figure 1A). The number of distinct plasmid sequences per genome (i.e. one representative of each, regardless of estimated copy number) ranged from 0-10 overall, with marine isolates typically having a higher number of plasmids per isolate (0-9 plasmids, median=4) than those from the human (0-10 plasmids, median=2) and animal (0-8 plasmids, median=2) niches. However, the differences were not significant (Kruskal-Wallis, p>0.05) (Figure 1B).

**Figure 1.**
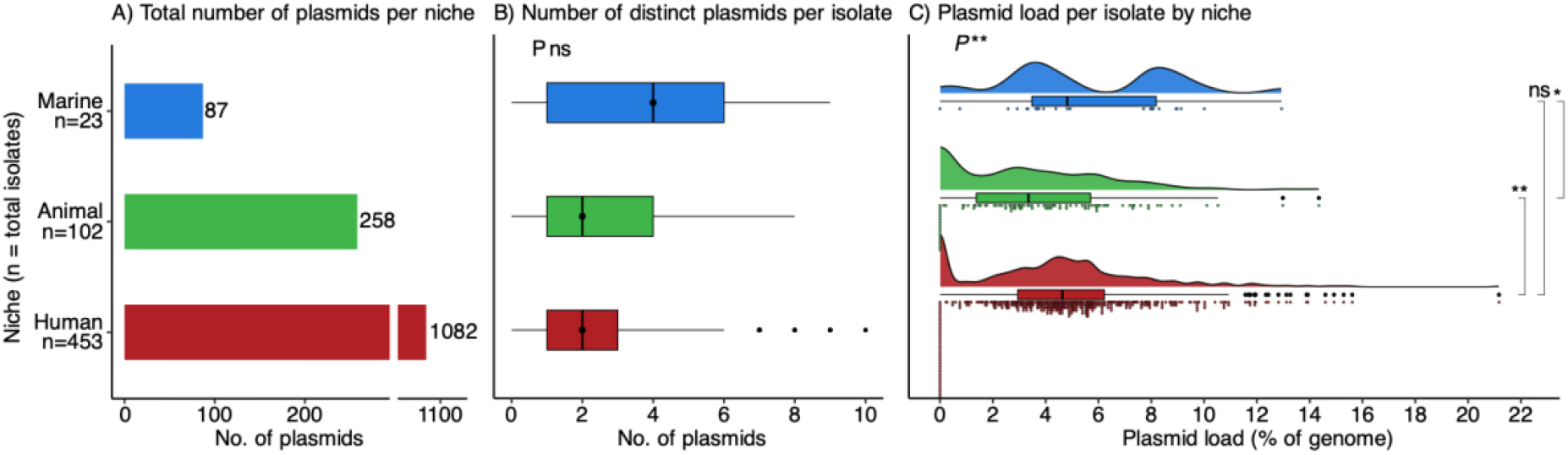
Plasmid distribution across niches. **A)** Total number of plasmids recovered per niche. **B)** Distribution of the number of distinct plasmid sequences per isolate within each niche, black vertical line indicates median value. **C)** Distribution of estimated isolate plasmid load within each niche. Statistical comparisons were performed using Kruskal-Wallis (overall) and Mann-Whitney (pairwise). Significance is denoted as follows: ^*^P<0.05, ^* *^P<0.01, ns P≥0.05.

While the number of distinct plasmids per isolate did not vary significantly by niche, the overall plasmid burden did. Plasmid load (i.e. percentage of each genome consisting of plasmid sequence when accounting for estimated copy number, see Methods) ranged from 0-21.2% (median 5.5%) overall. It was significantly lower in the animal niche (0-14.3%, median 3.4%) compared to the human (0-21.2%, median 4.6%) (p<0.01) and marine (0-12.9%, median 4.8%) (p<0.05) niches (Figure 1C).

Most plasmids were found in *K. pneumoniae* (87.0%, n=1241/1427 plasmids among 447 isolates), followed by *K. variicola* (10.2%, n=146/1427 plasmids among 55 isolates), *K. quasipneumoniae* subsp. *similipneumoniae* and *quasipneumoniae* (1.1% and n=16/1427 plasmids each among 7 and 4 isolates, respectively) (Figure S1A), and a single *K. quasivariicola* isolate harboured 0.6% of all plasmids (n=8/1427). While both the number of distinct plasmids per isolate (Figure S1B) and estimated plasmid load differed between species (Figure S1C), the only statistically significant difference was the higher plasmid load in *K. pneumoniae* (0-21.2%, median 4.6%) versus *K. quasipneumoniae* subsp. *similipneumoniae* (0-5.1%, median 2.1%, p<0.05).

### Plasmids were diverse within and across niches

The 1415 circularised plasmids were used to characterise, cluster, and compare plasmids across the niches. Among the circularised plasmids, a total of 2013 plasmid replicon markers (69 distinct markers) were identified (Table S1). All but one plasmid contained 0-4 replicon markers per plasmid (median=1) with slight variation across niches (Figure 2A). The majority of plasmids harbored a single replicon marker (n=826/1415, 58.3%), the most prevalent were ColRNAI_rep_cluster_1987 in both the human and marine niches (203 and 20 plasmids, respectively), and Col(MG828) and rep_cluster_2401 in the animal niche (in 27 plasmids each). In all niches, the second most common replicon on single-replicon plasmids was IncFIB (135, 26, and 5 plasmids across human, animal and marine niches, respectively). Plasmids harboring two replicon markers (14.8% n=210/1415) comprised the second largest portion of plasmids, with IncFIB/IncFII being the most frequent combination in the human and animal niches (62 and 23 plasmids, respectively) and IncFIB/rep_cluster_1418 in the marine niche (3 plasmids). The remaining plasmids harboured zero, three, or four replicon markers (n=131, n=221, n=26, respectively).

**Figure 2.**
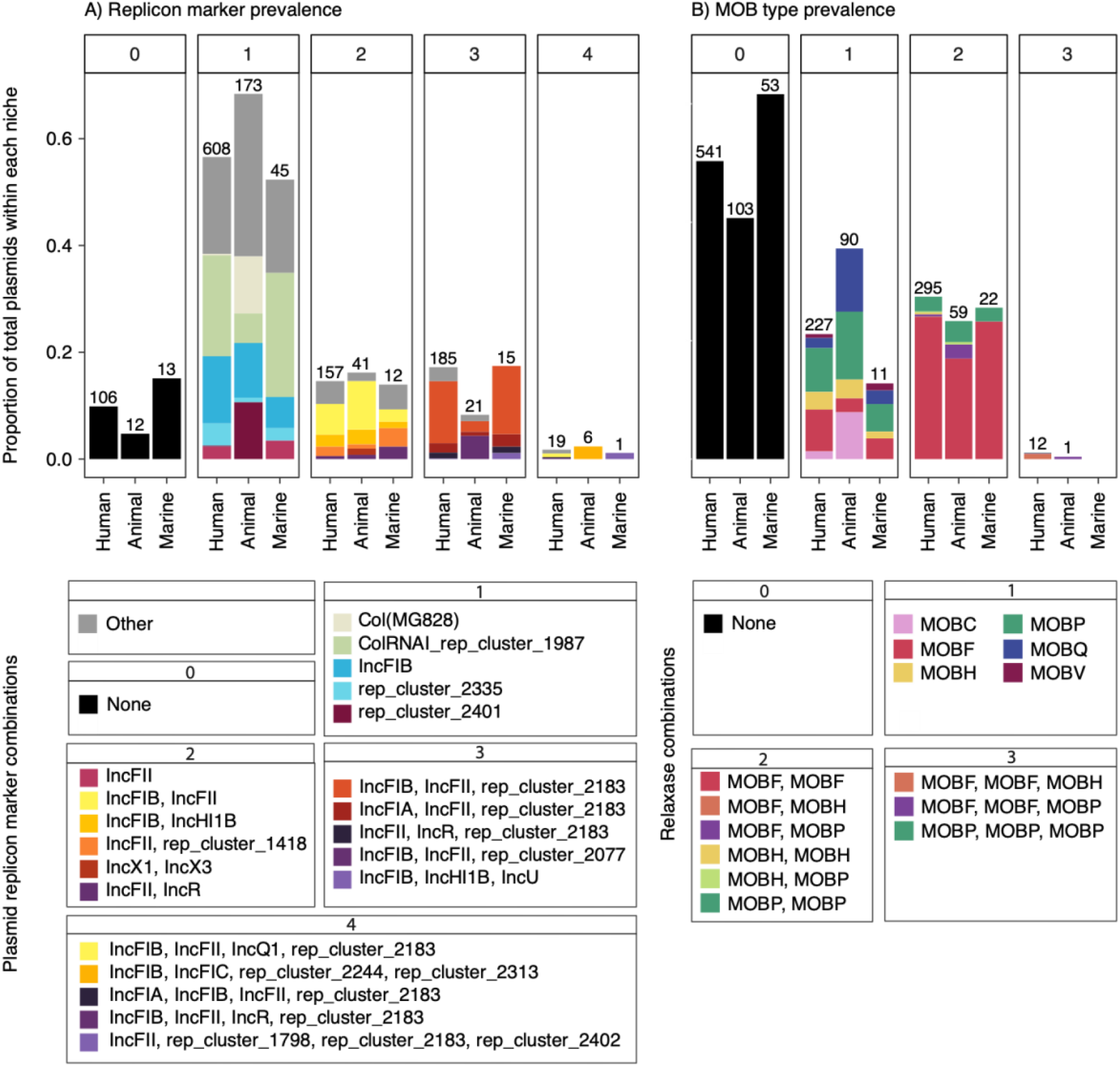
Replicon and MOB type distribution across niches. **A)** Replicon marker prevalence by niche. Facet numbers indicate the number of markers per plasmid. Bars indicate the proportion of plasmids within each niche harbouring different replicon markers/combinations; total number of plasmids with specified number of replicon markers per niche above each bar. The three most frequent marker types for each niche/replicon number combination are shown, the rest are categorised as “Other”. **B)** MOB type prevalence by niche. Facet numbers indicate the number of relaxases per plasmid. Bars indicate the proportion of plasmids within each niche harbouring different relaxase combinations corresponding to the number of relaxases per plasmid; total number of plasmids with specified number of relaxases per niche above each bar.

Most plasmids (59.2%, 838 /1415) were predicted to be transmissible (n=486 conjugative, n=352 mobilisable), while 40.8% (n=578 /1415) were predicted to be non-mobilisable. In total, 1126 relaxases belonging to MOBC, MOBF, MOBH, MOBP, MOBQ, and MOBV types were identified. Plasmids carried 0-3 relaxases (median=1) in various combinations. Nearly half of plasmids did not harbour a known relaxase (49.2%, 697 /1415), of these, 17.7% (n=119/697) were predicted to be mobilisable based on the presence of an origin of transfer. Almost a quarter of all plasmids harboured a single relaxase (n=328/1415; 23.2%) and corresponded to the largest portion of plasmids from the animal niche (35.6%, 90/253); they most frequently encoded MOBP in all niches. Plasmids carrying two relaxases were the largest groups in the human and marine niches (27.4% and 25.6%, respectively); the most common pairings were MOBF/MOBF followed by MOBP/MOBP in all niches (Figure 2B).

One plasmid harboured more than four replicon markers; this was likely misassembled as it was much larger than other plasmids assigned to the same cluster (>149kbp) and contained exact combinations of replicon and relaxase markers observed in other plasmids roughly half its size assigned to the same grouping in the cluster analysis (below); it was therefore reclassified as a hub plasmid in the cluster analysis, and excluded from replicon marker and relaxase visualisations and calculations below.

### Clinically relevant plasmid-borne features varied across niches

Of the 514 genomes that carried plasmids, 28.6% (n=147/514) carried ≥1 plasmid encoding AMR genes, corresponding to a total of 182 AMR-encoding plasmids (12.9% of all plasmids). Virulence genes or loci (*iuc, iro, rmpADC, rmpA2*, or *ybt*, excluding truncated or incomplete hits) were detected on plasmids in 14.8% (n=76/514) of plasmid-containing genomes, across 76 plasmids (5.4%). HMR genes were more frequently observed, found in 66.0% (339/514) of genomes and encoded by 384 plasmids (27.1%). A subset of genomes (28.0%, n=144/514) harboured multiple plasmid-encoded clinically relevant traits, with features either carried on the same or on separate plasmids. In total, 118 (23.0%) genomes encoded AMR and HMR genes; 11 (2.1%) carried HMR and other virulence genes; four genomes (0.7%) carried AMR and virulence genes; and 11 genomes (2.1%) harboured genes conferring AMR, HMR, and other virulence factors (Figure S2).

Many genomes with several clinically relevant features could be explained by the presence of single plasmids harbouring these traits. The most frequent combination was AMR and HMR genes, co-encoded on 7.7% (109 /1415) of plasmids. HMR and other virulence genes were detected on 1.2% (17 /1415) of plasmids. Five plasmids (3.5%) carried both AMR and virulence genes, three of which also encoded HMR (Figure 3). Of these, only one plasmid (belonging to an SL15 isolate from human blood) encoded an ESBL (*bla*_CTX-M-15_) together with the *iuc1* locus, and mercury resistance genes (*merACDEPRT*) (Figure S3).

**Figure 3.**
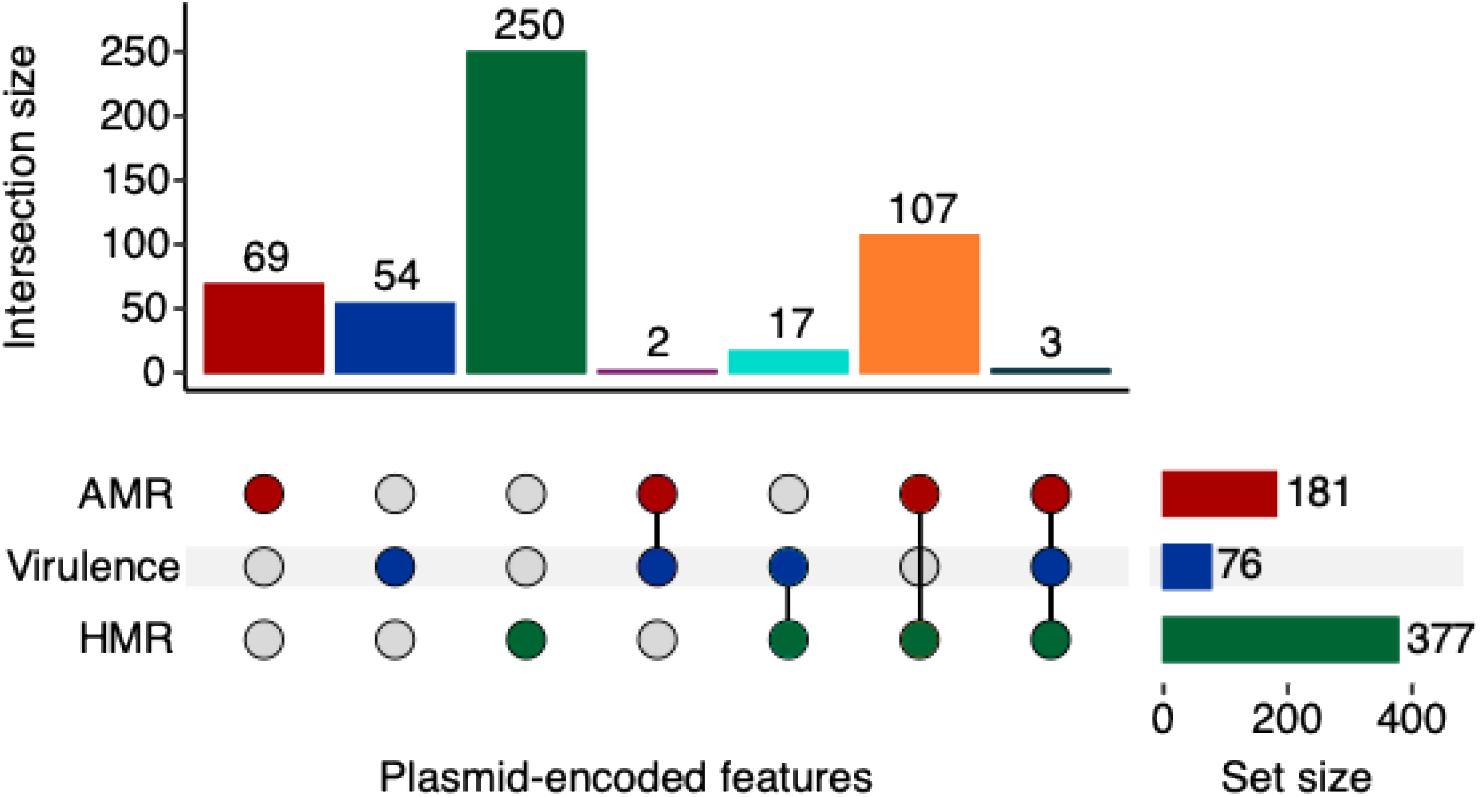
Overlap of clinically relevant features across plasmids. Upset plot showing the overlap of clinically relevant features harbored across n=508 /1415 plasmids. AMR genes in red, virulence factors in blue, and HMR genes in green.

The convergent AMR and virulence plasmids (n=5) were only observed among human infection isolates belonging to SL15 (n=2), SL641 (n=1), SL846 (n=1), and SL881 (n=1), and harboured at least an IncFIB replicon. Four were predicted to be conjugative and encoded MOBF (n=2), MOBH (n=1), or MOBF/MOBP (n=1) relaxases, and one plasmid lacked a relaxase and was predicted to be non-mobilisable. Of the 182 plasmids carrying AMR genes, 114 conferred MDR. Plasmids encoding AMR were present in all niches but primarily associated with the human niche (164/1077, 12/253, and 6/86 plasmids in the human, animal, and marine niches, respectively). The *iuc*3, *iuc*5, and *iro*5 virulence loci were enriched in the animal niche (p<0.05), with *iuc3* specifically associated with pigs, as previously reported (25,35) and *iuc5+iro5* loci linked to a previously reported SL290 clonal expansion in turkeys (19), while *iuc1* was enriched in the human niche (p<0.05).

Overall we observed high plasmid diversity in this collection, with sequence lengths ranging 1.2-424 kbp (median=57 kbp), however AMR genes were present across plasmids of a broad size range, 2.9-353 kbp (median=157 kbp), while virulence factors were typically encoded on 120-250 kbp (median=177 kbp) plasmids (Figure S4).

AMR genes and/or virulence factors were observed on plasmids with a variety of replicon markers and MOB types (34 replicons, 5 MOB types). We observed correlation of several AMR and virulence genes with particular replicon or MOB types (Figure S5). Among the ESBL-encoding genes, *bla*_CTX-M_ variants (particularly *bla*_CTX-M-3_, *bla*_CTX-M-14_, and *bla*_CTX-M-15_) were most commonly found on MDR plasmids harbouring IncFIB and IncFII replicons and ≥1 MOBF relaxase. The siderophore-encoding loci (*iuc, iro*) and capsule regulators (*rmpA, rmpA2*) were also most frequently found on plasmids harbouring at least one IncFIB and one IncFII replicon and ≥1 MOBF relaxase. While IncFIB/IncFII and MOBF/MOBF plasmids were also common amongst plasmids lacking AMR genes and/or virulence loci, the majority of these plasmids harboured a ColRNAI_rep_cluster_1987 replicon and lacked a relaxase.

### Transposable elements (TEs) and plasmid mobility differed by niche and gene content

Both AMR genes and virulence factors primarily resided on conjugative plasmids containing 0-42 (median=12) TEs on AMR plasmids and 2-35 (median=6) TEs on virulence plasmids, with the exception of *iuc*1-encoding plasmids, which harboured more TEs on average compared to other virulence plasmids (12-35, median=15) and most were putatively non-mobilisable (n=11/15). Insertion sequences (ISs) were the dominant TE (Figure S6A-D), accounting for 98.0% of TEs. Plasmids harbouring AMR genes exhibited significantly higher counts of IS elements (0-42, median=14) compared to those without AMR (0-47, median=0) (p<0.001) (Figure S6B), implying a potential link between IS count and AMR in plasmids.

TE profiles also varied by ecological niche (Table S1). Plasmids from both human and marine sources contained significantly more IS elements (0-47, median=7; 0-37, median=2, respectively) compared to those from the animal niche (0-35, median=0) (p<0.01 and p<0.001, respectively). However, miniature inverted-repeat transposable elements (MITEs) were significantly enriched in the animal niche (6.30%, n=16/254 plasmids) compared to the human niche (1.49%, n=16/1077 plasmids) (p<0.001). Unit transposons were more frequent in plasmids from the human niche (9.75%, n=105/1077) compared to the animal niche (3.15%, n=8/254) (p<0.001) but not the marine niche (6.06%, n=6/99) (p>0.05).

### KpSC plasmid clustering showed high diversity

Clustering the circularised plasmids (n=1415) with Pling (see details in Methods) resulted in 49 communities (i.e. plasmids within the containment distance threshold of 0.5). These were broken down further into 130 subcommunities (plasmids within 4 rearrangements of each other, hereafter referred to as “clusters”), 524 singletons, and 13 hub plasmids (highly connected plasmids that lead to overclustering). The plasmid collection exhibited considerable variation in size and content. The 49 communities ranged in size from 2-1120 plasmids per community (median=3), encompassing 1363 plasmids (Figure 4). The largest community accounted for the vast majority of plasmids overall (79.1%, n=1119 /1415), including 81.9% (n=429/524) of all singletons, and all hub plasmids. Excluding hub plasmids, a total of 878 plasmids grouped into 130 distinct clusters ranging in size from 2-117 plasmids per cluster (median=16). The 524 singleton plasmids accounted for 37.0% of the overall collection with a notable portion of those found in each of the human, animal, and marine niches (40.5%, 22.9%, and 34.9% of each niche, respectively), demonstrating substantial plasmidome diversity. Pling detected ten hub plasmids and an additional three were identified upon inspection of the network clusters: ten were found in the human niche, of which two encoded ESBLs (*bla*_CTX-M-15_ and *bla*_SHV-2_); three hubs were from the marine niche, none of which encoded AMR or virulence. All hub plasmids were excluded from network visualisation (Figure 4; see Figure S7 for a complete network including hubs and singletons).

**Figure 4.**
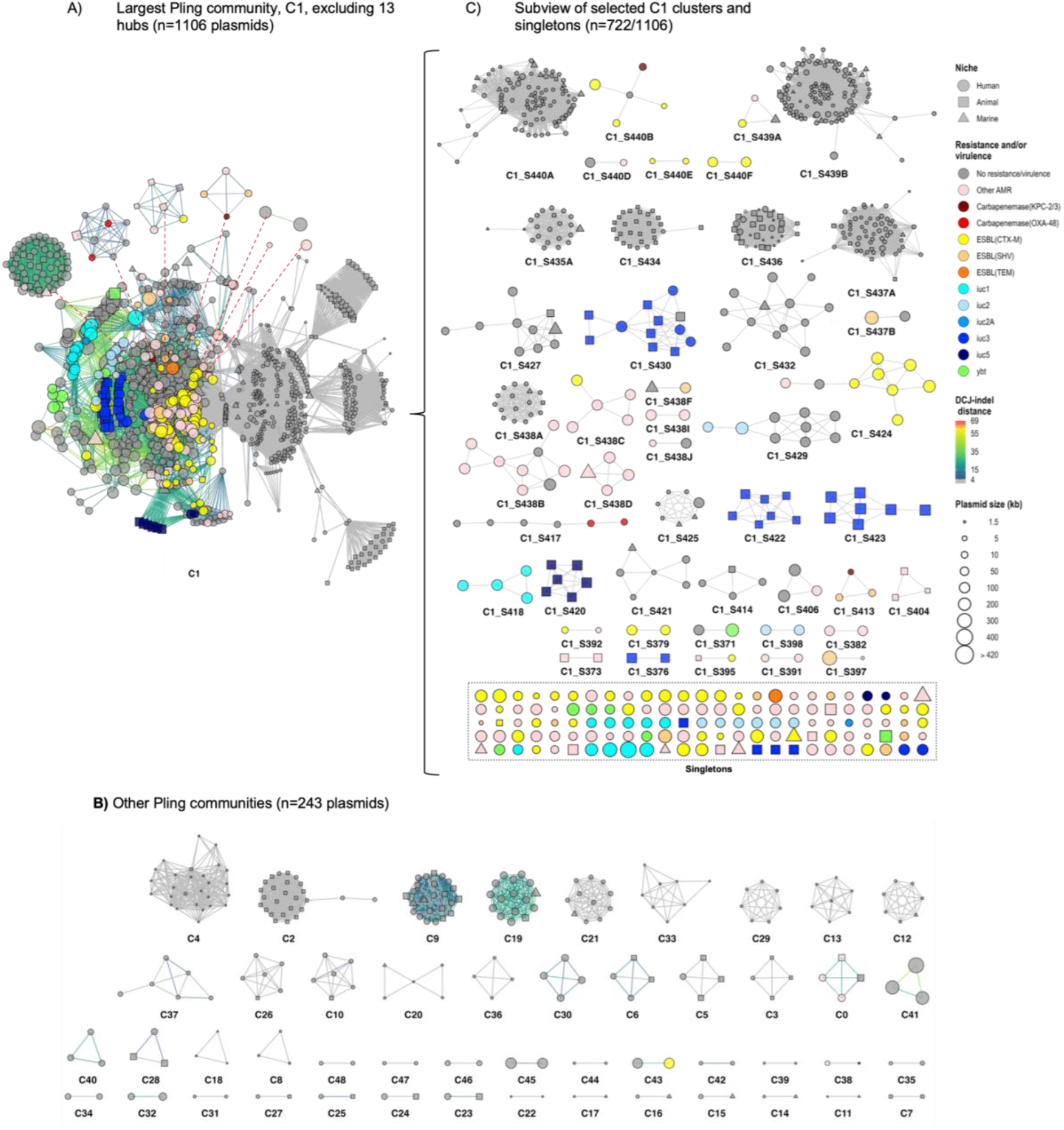
Pling network graph of n=1415 circularised plasmids. **A)** and **B)** Plasmids belonging to n=49 Pling communities (n=1350 plasmids), excluding 13 hub plasmids; red dotted lines indicate cluster connections that would have been made via hub plasmids. **C)** Subview of selected clusters and singletons within community C1. Shown are clusters that (i) contain ≥1 AMR and/or virulence plasmid, or (ii) comprise ≥4 plasmids derived from ≥2 niches, and (iii) singletons that encode ≥1 AMR gene and/or virulence factor (shown in box). Node shape indicates isolate niche, node size indicates plasmid length in kbp, node color represents clinically relevant plasmid-encoded features. Edge color indicates DCJ-indel distance between plasmids within the same community, where grey indicates ≤4 rearrangements between sequences. Not shown: clusters containing <4 plasmids (n=85 clusters), hub plasmids (n=13), and singletons lacking clinically relevant genes (n=391 plasmids). For the complete Pling network, including hub plasmids and all singletons, see Figure S7.

Excluding the 13 hub plasmids, AMR and/or virulence genes were detected in 37 clusters and 133 singletons (n=246 /1415 plasmids); 28 clusters and 101 singletons contained at least one AMR plasmid (n=172 /1415 plasmids); nine clusters and 31 singletons contained plasmids encoding virulence factors (n=71 /1415 plasmids). Five singleton plasmids contained both AMR and virulence genes. All were found in human infection isolates of various SLs (SL641, n=1; SL846, n=1; SL881, n=1; and SL15, n=2). Four of the five plasmids encoded resistance to aminoglycosides; the fifth, found in SL881, encoded *iuc3* and tetracycline resistance. The plasmid from SL846 *iro5*, while the plasmid from SL641 encoded both *iuc5* and *iro5*. The two plasmids from SL15 isolates encoded *iuc1*, and one of these plasmids also harboured an *bla*_CTX-M-15_ and *bla*_SHV-5_.

### Within-cluster diversity

There were 9711 annotated genes among the 1415 plasmids. Within each cluster, we compared pairwise Jaccard distances, and categorised genes as core (present in ≥90% of plasmids) or accessory (<90% of plasmids) (Figure 5).

**Figure 5.**
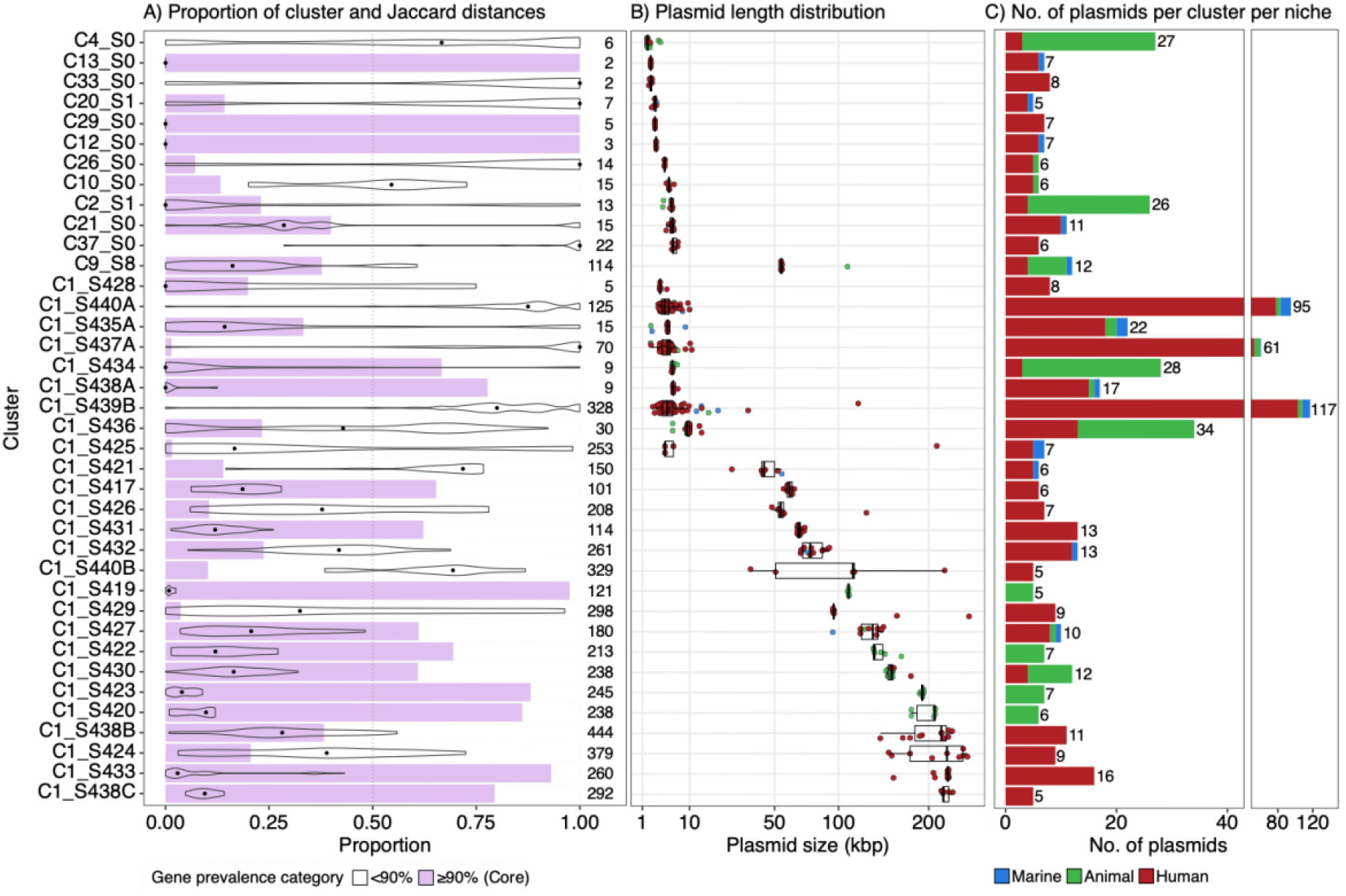
Within-cluster diversity. **A)** Proportion of genes in selected clusters that were found in ≥90% of cluster plasmids (i.e. core genes relative to cluster, purple), and distribution of within-cluster pairwise Jaccard distances between plasmids; sum of the total number of Bakta-annotated genes per cluster next to each bar. **B)** Distribution of plasmid lengths within each cluster (kbp), colored according to isolate niche. **C)** Number of plasmids per cluster from each of the human (red), animal (green), and marine (blue) niches. Not shown: clusters with <5 plasmids (n=92 clusters).

There were 38 clusters containing ≥5 plasmids, which accounted for 53.1% of plasmids (n=752 /1415). Five of these clusters lacked any core genes. These included four diverse clusters with plasmids from all three niches, including the two largest clusters (n=117 and n=95 plasmids). Interestingly the fifth cluster lacking core genes contained plasmids from the human niche (n=6 plasmids) within a narrow length range (5.6-6.9 kbp), suggesting that some plasmid clusters may have been defined by accessory gene content rather than shared core features.

Of 130 clusters (excluding hub plasmids), 47 (36.2%) included plasmids originating from multiple niches (15 human-animal, 22 human-marine, 3 animal-marine, and 7 from all three niches), accounting for 42.9% (n=607 /1415) of plasmids overall (see Figure 4). Within our collection, some SLs were more promiscuous than others, and some clusters tended to represent more diverse SLs than others. Among the 255 SLs, the number of plasmid clusters ranged from 1 to 53 (median=3). Plasmid clusters were observed across 1 to 63 SLs, although the majority were confined to a single SL (median=1) (Figure S8).

Some clusters appeared highly conserved, for example, clusters C1_S376, C1_S422, C1_S423 and C1_S430 comprised 28 *iuc*3-encoding plasmids (one of which was a truncated variant) predominantly from the animal niche (n=23 pig, n=1 dog), though C1_S430 also contained plasmids from the human niche (n=3 infection, n=1 carriage). Plasmids in these clusters were of similar length (134-202 kbp, median=156 kbp) and all harboured MOBF, MOBF relaxases as well as IncFIB/IncFII replicon markers, however they were distributed across diverse SLs, with the only overlaps occurring in SL35 (n=1 pig, n=1 human infection) and SL37 (n=3 pig, n=1 dog). Another conserved cluster, C1_S427, contained ten plasmids from all three niches (n=8 human, n=1 animal, n=1 marine), all lacking AMR or virulence genes and predicted non-mobilisable with a single IncFIB(K) replicon. Interestingly, the plasmid from the animal niche (turkey) carried the same HMR operons (*arsABCDR, pcoABCDRS, silABCERS*) as six of the plasmids from the human niche (five from human infections, one from carriage). While plasmids from the turkey isolate and two of the human infection isolates were harboured by the same SL (SL152), the remaining four plasmids from human infection and one carriage isolate were each harbored by distinct SLs.

Other clusters appeared far more diverse at first glance, such as C1_S438 and C1_S440, however, further inspection and removal of three additional hub plasmids resulted in slightly more homogenous clusters. Clusters C1_S438A-B initially belonged to the same group (C1_S438) but were further divided based on plasmid length. C1_S438B was a highly conserved cluster consisting of 17 5.9-6.9 kbp plasmids from three niches (n=15 human, n=1 animal, n=1 marine), and all encoding a single ColRNAI_rep_cluster_1987 replicon and no relaxases, AMR or virulence genes. However, C1_S438A remained a diverse group of 62 plasmids (n=56 human, n=2 animal, n=4 marine) ranging in length from 87.1-282.6 kbp: all harbored three replicon markers, one of which was IncFII (followed in declining frequency by IncFIB, IncFIA, IncQ1, rep_cluster2183, and/or rep_cluster_1418). C1_S440 was also divided into smaller clusters. C1_S440A contained 95 plasmids spanning all three niches, all of which were <11 kbp and lacked a relaxase, and most of which were predicted to be non-mobilisable with a ColRNAI_rep_cluster_1987 replicon marker. While the plasmids assigned to clusters C1_S440B-F were considerably larger (32-424 kbp) and represented all three niches, AMR genes (*bla*_CTX-M-15_, *bla*_KPC-2_, genes encoding resistance to aminoglycosides, etc.) were only present in plasmids from human sources, reflective of plasmids being shaped by their environment.

### The Norwegian KpSC plasmids are representative in a global context

To contextualise the Norwegian KpSC plasmidome within a broader genomic landscape, we compared our collection to 8656 circularised plasmids from publicly available KpSC genomes downloaded from the NCBI RefSeq (see details and inclusion criteria in Methods), collected from 65 countries in six world regions between 2015 and 2024 (Figure 6A, Table S2). Nearly half of the plasmids were from isolates collected in Asia (n=4960), followed by Europe (n=3016, including plasmids from this study), North America (n=1246), Oceania (n=467), South America (n=272), and Africa (n=111), resulting in a total of n=10072 plasmids in the global dataset (Figure 6A). To identify overlap between our collection and the global dataset, the European region was divided into: ‘Europe’ (n=1600, including 6 Norwegian plasmids not from this study), and ‘Norway’ (plasmids in this study, n=1415). Clustering with Pling resulted in 90 communities, 614 clusters, 1544 singletons (15.3%), and 363 hub plasmids (3.6%) (Figure S9). Interestingly, plasmids from our Norwegian collection were broadly dispersed throughout the global network: they were present in 66.1% of multi-region communities (n=37/56), and 63.2% of multi-region clusters (n=172/272). This distribution indicates broad overlap between plasmids in this dataset and the global KpSC plasmid landscape (Figure 6B). Nine communities (10.0%), 42 clusters (6.1%) and 396 singletons (23.9%) consisted exclusively of plasmids from this study. As with the Pling output of our dataset, the global dataset was dominated by a single large community that accounted for the vast majority of plasmids in the dataset (92.0%, n=9265/10072). Most communities (62.2%, n=56/90) represented >1 world region. Excluding hub plasmids, 39.4% of clusters (n=272/690) contained plasmids from >1 world region (Figure 6B). Of these, eight clusters contained plasmids from all six regions and accounted for 22.4% of all plasmids (n=2255/10072).

**Figure 6.**
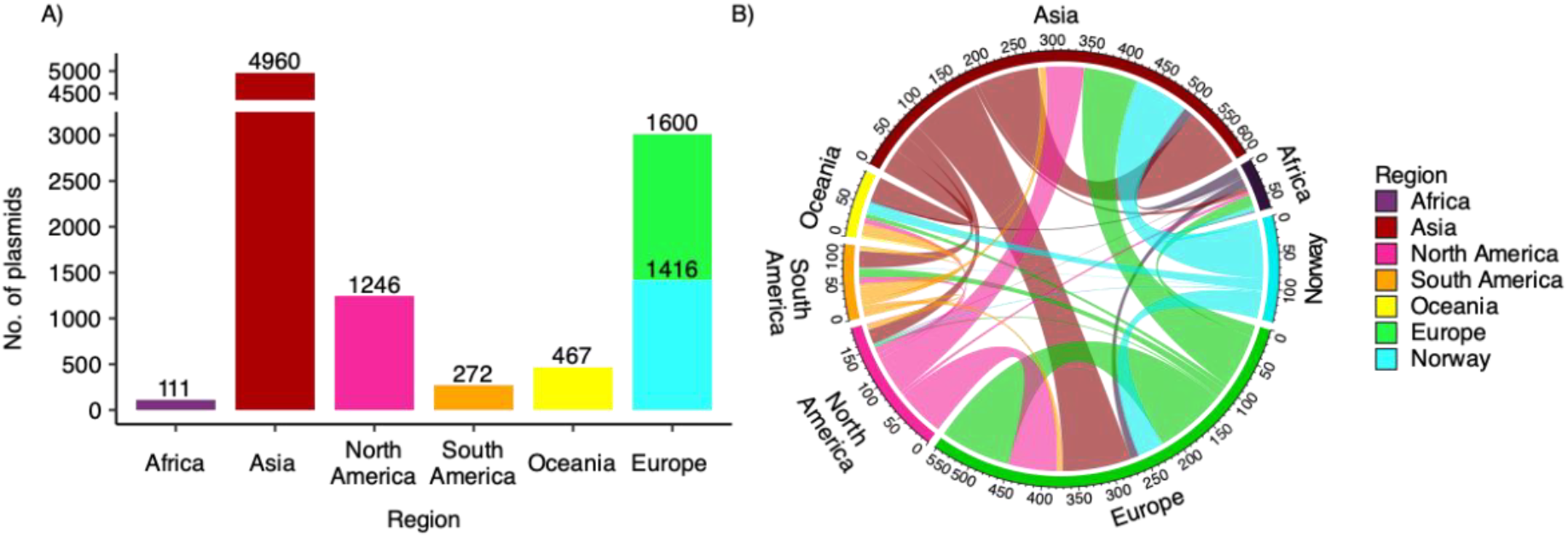
Distribution of global KpSC plasmids. **A)** Barplots indicate the total number of plasmids included in global clustering analysis from each world region. **B)** Chord diagram of plasmids assigned to the same clusters and found in multiple geographic locations. See Fig S10 for alternative world map views.

Excluding hub plasmids, a total of 328 clusters and 567 singletons contained plasmids harbouring either AMR and/or virulence genes. Of these, 282 clusters and 461 singletons contained plasmids encoding AMR, 38 clusters and 59 singletons contained virulence plasmids, and 32 clusters and 47 singletons contained plasmids encoding both AMR genes and virulence factors. Among hub plasmids (n=363), 61.7% (n=224) carried AMR genes, 0.8% (n=3) encoded virulence factors, and 8.5% (n=31) encoded both features.

Of the 282 clusters containing AMR plasmids, 42.6% (n=120) spanned >1 global region, 34 of which also contained AMR plasmids from our dataset. However, many plasmids from this dataset that did not encode AMR genes were also found in global clusters containing AMR plasmids from other global regions. Most of the 38 clusters containing virulence-only plasmids spanned >1 global region (52.6%, n=20), and 14 of those also contained virulence plasmids from this dataset, with each virulence locus represented in ≥1 cluster. Of the 31 clusters containing plasmids that encoded both AMR and virulence, 54.8% (n=17) spanned 2-6 global regions (2 regions n=5 clusters; 3 regions n=5; 4 regions n=1; 5 regions n=3; 6 regions n=3), of which only one cluster contained plasmids from our dataset.

## Discussion

This study provides a comprehensive analysis of the plasmid diversity and associated AMR and virulence factors in KpSC isolates across three ecological niches in Norway. Our findings demonstrate the extensive diversity and adaptability of KpSC plasmids, underlining their role in AMR and virulence transmission within and across sectors.

The variation in plasmid load across niches, with human and marine isolates harbouring significantly greater plasmid loads than animal isolates, suggests that these environments may impose distinct selective pressures or support ecological conditions conducive to plasmid retention and acquisition. Additionally, as KpSC marine isolates were predominantly detected near coastlines rather than in open water (18), it is likely that isolates and plasmids from human and marine sources originated from similar backgrounds as a result of wastewater and/or terrestrial runoff, or via human consumption of seafood (36,37). While plasmid diversity was widespread across KpSC species with only slight variation in plasmid burden between groups, ecological niche appeared to play a more significant role in shaping plasmid contents.

Niche-specific patterns were particularly evident in the distribution of AMR and virulence plasmids. The human niche, as expected, had a significantly higher prevalence of AMR plasmids, likely reflecting the impact of human antibiotic use (2,3,9,38–41) relative to the lower use of antibiotics in the agricultural sector in our settening(42,43). In contrast, the animal niche was enriched for virulence-encoding plasmids, specifically those carrying *iuc3* and *iuc5*. The *iuc3* locus has previously been linked to pig-associated lineages (24,35), while *iuc5* was associated with a clonal expansion in turkeys (25). Interestingly, these were predominantly found in non-HV clones (1), highlighting the potential role of animal reservoirs in the evolution of virulent KpSC strains that may cross into human populations. Conversely, *iuc1* and *iuc2* plasmids were found exclusively in the human niche and were associated with globally recognised HV clones and even some MDR lineages (1). The distribution of clinically relevant plasmids across both high-risk MDR clones and other SLs further demonstrates the capacity of KpSC to acquire and retain plasmids across diverse clonal backgrounds, and suggests plasmid-mediated gene flow is not restricted solely to high-risk clonal groups.

Clinically relevant genes were frequently co-carried with well-characterised replicon markers across all niches. In particular, plasmids encoding *bla*_CTX-M-3_, *bla*_CTX-M-14_, and *bla*_CTX-M-15_ were consistently associated with IncFIB and IncFII, while siderophore loci (*iro, iuc*, excluding *iuc2A*) and capsule regulators (*rmpA, rmpA2*) were linked to IncFIB(K) and IncFIB. Although most AMR-encoding plasmids were predicted to be conjugative or mobilisable, several *iuc1*-carrying plasmids (in the Norwegian and global plasmid collections) lacked conjugation machinery and were predicted to be non-mobilisable, consistent with previous findings by Lam et al. (10), suggesting alternative dissemination routes such as clonal expansion or mobilisation by co-resident plasmids. The association of MOBP and MOBF relaxases with plasmids encoding ESBLs and siderophores further supports their role in mobilising clinically relevant plasmids across ecological and genomic contexts (14).

The TE burden on AMR plasmids compared to non-AMR counterparts suggests that TEs play a disproportionate role in AMR gene acquisition and structural rearrangement. Many of the 120-250 kbp plasmids encoding both AMR and virulence genes had multiple IS elements, implicating TEs in facilitating convergence events. Together, these findings support a model in which replicon type, MOB machinery, and TE content shape both the composition and dissemination potential of clinically important plasmids.

A significant proportion (37.0%) of the Norwegian collection consisted of singletons, reflecting a highly dynamic plasmidome. Additionally, many plasmids (42.9%) were assigned to multi-niche clusters, or shared considerable sequence content with global counterparts, indicating that many plasmids are capable of persisting and spreading across broad ecological and geographical ranges. Despite Norway’s relatively low antimicrobial usage across multiple sectors (42,44), Norwegian plasmids showed substantial overlap with the global KpSC plasmidome, with Norwegian plasmids present in over half of the global Pling clusters. While many of the Norwegian plasmids lacked AMR genes, they frequently clustered with globally distributed AMR plasmids. This suggests that gene-acquisition-ready plasmids are already circulating in low-AMR settings and highlights the importance of plasmid backbone surveillance. These patterns reinforce the notion that plasmids act as vehicles of gene flow across both microbial and environmental boundaries, and underscore the global interconnectedness of plasmid evolution and the critical need for One Health-informed genomic surveillance, including plasmids.

Although we did not aim to infer direct plasmid transmission events between niches in our dataset or across geographies with the global dataset, Pling clusters of structurally similar plasmids, paired with other relevant genetic markers (i.e. combinations of replicon markers, relaxases, AMR genes and virulence loci) suggest that dissemination occurs both across niches and country borders. We have previously shown that there were relatively few strain-sharing events across ecological niches, compared to within-niche strain-sharing (4). Our findings here suggest that plasmid-sharing is considerably more common across niches than strain-sharing. However, it remains unclear if these plasmids have evolved from a common ancestral source, or have disseminated from one niche to another via intermediaries.

There are several strengths and limitations in this study. While this study includes a large, hybrid-assembled dataset of KpSC isolates spanning three ecological niches and two decades across Norway, the collection was skewed toward human isolates due to availability, and plasmid-rich isolates were prioritised for long-read sequencing. However, approximately 20% of the collected samples from each niche were long-read sequenced, ensuring proportional representation. The reliance on isolates collected primarily from human sources likely influenced the observed distribution of plasmid types and associated genetic elements. Moreover, while our global comparison included high-quality complete genomes, it remains constrained by public database biases, particularly overrepresentation of clinical isolates, imbalanced representation of world regions, and underreporting of ecological metadata.

Nonetheless, this study represents one of the largest and most comprehensively analysed plasmid datasets from a national One Health perspective. The use of hybrid assembly and plasmid clustering allowed us to resolve full plasmid structures, enabling comparative analyses at a high resolution. The surprising degree of overlap between our plasmid collection and global plasmid clusters highlights the utility of such national collections in global AMR surveillance efforts and underscores the global relevance of plasmid populations from low-AMR settings.

Future work should incorporate more extensive sampling from underrepresented hosts, environments, and geographical locations, along with longitudinal monitoring to track plasmid evolution. Further investigations into the functional impact of plasmid diversity - particularly of small, cryptic plasmids and phage-plasmids - are needed, as these may play underappreciated roles in HGT. Experimental validation of plasmid fitness effects in different contexts would also illuminate the mechanisms underlying their persistence and spread.

In conclusion, the plasmidome of our KpSC collection is diverse, globally representative, and shaped by ecological niche. Similar plasmid backbones capable of acquiring and transmitting clinically significant traits are already circulating across human, animal, and marine niches in Norway. These findings underscore the importance of continued plasmid-level surveillance and integrated One Health strategies to prevent the spread of AMR and virulence traits.

## Data availability

The Illumina and Oxford Nanopore Technologies reads for all isolates are available under project PRJEB74192 on the European Nucleotide Archive. The hybrid-assembled genomes have been deposited in GenBank. See Table S1 for metadata and genotyping results of the Norwegian dataset, and Table S2 for accessions and genotyping results of the global dataset.

Acknowledgements

We wish to thank the NOR-KLEB-NET and KLEB-GAP partners, the Institute of Marine Research in Norway, the Norwegian Veterinary Institute, the Norwegian Centre for Detection of Antimicrobial Resistance, the Tromsø7 population survey, and the Norwegian Surveillance Program on Resistant Microbes (NORM)/NORM-VET programs for their efforts in collecting and/or providing samples of KpSC, that were whole-genome sequenced at the Department of Medical Microbiology, Stavanger University Hospital, Norway. We would also like to thank Zamin Iqbal and Daria Frolova for constructive discussions and valuable input on Pling analysis.

## Author Contributions

**Conceptualisation**: MAW, MAKH, ØS, AS, MMCL, IHL. **Methodology**: MAW, MAKH, JH, ØS, MMCL. **Software**: MAW, MAKH, JH, MMCL. **Formal analysis**: MAW, MAKH. **Investigation**: RJB, EB. **Resources**: HK, AF, BTL, NPM, NR, ØS, MS, AS, IHL. **Data Curation**: MAW, MAKH. **Writing - Original Draft**: MAW, MAKH. **Visualisation**: MAW, MAKH. **Supervision**: ØS, MMCL, IHL. **Project administration**: MAW, IHL. **Funding acquisition**: MAKH, BTL, MS, ØS, AS, IHL. **Writing - Review & Editing:** MAW, MAKH, HK, RJB, AF, NR, NPM, BTL, MS, JH, EB, ØS, AS, MMCL, IHL.

## Funding

This study was supported by grants from the Trond Mohn Foundation (TMF2019TMT03) and from the Western Norway Regional Health Authority (F-12508 to MAKH).

## Conflicts of interest

The authors declare that there are no conflicts of interest.

## Ethics declarations

The North Norwegian Regional Committee for Medical and Health Research Ethics gave approval to the collection of isolates from human fecal carriage (REC North reference: 2016/1788). The Regional Committee for Medical and Health Research Ethics in Western Norway gave its approval to the two studies that collected isolates of human infections (REC West reference 2017/1185 and 2016/1093). No additional ethical approvals were needed to use the human fecal carriage and infection KpSC genomes for this investigation as we did not gather any more samples or data. The genomes used in the three published studies are accessible to the general public (in BioProjects: PRJEB42350, PRJEB48268 and PRJEB27256). The study complied with the Helsinki Declaration’s guidelines.

### Abbreviations

AMR: antimicrobial resistance
cgMLST: core genome multilocus sequence type
DCJ-indel: Double Cut and Join insertions and deletions
ESBL: extended-spectrum β-lactamase
HGT: horizontal gene transfer
HMR: heavy metal resistance
HV: hypervirulent
Indel: insertion-deletion
IS: insertion sequence
KpSC: *Klebsiella pneumoniae* species complex
LIN: life identification number
MITE: miniature inverted repeat element
MDR: multidrug resistant
ONT: Oxford Nanopore Technologies
SL: sublineage
ST: sequence type
TE: transposable element

## Notes

### Competing Interest Statement

The authors have declared no competing interest.

