## Supplementary methods for "A One Health study of *Klebsiella pneumoniae* species complex plasmids shows a highly diverse and ecologically adaptable plasmidome"

### Sample selection for long-read sequencing

The isolates selected for long-read sequencing were part of a larger dataset (n=3255) that had been previously short-read sequenced in several published studies (1–10). To generate a representative subset of *Klebsiella pneumoniae* species complex (KpSC) isolates for hybrid genome assembly, we applied a multi-step selection process from the initial collection of 3,255 isolates. Our aims were to ensure a representative selection of the dataset, capture clinically relevant antimicrobial resistance and/or virulence genes, and broad representation of the plasmid diversity, while optimizing sequencing efficiency.

The initial selection of 359 genomes prioritized representativeness of the overall collection after analysis with Bakta v1.8.1 with database v5.0 (11), Panaroo v1.3.3 (12), and a long-read selection pipeline (https://gitlab.com/sirarredondo/long_read_selection), as well as ensuring each sublineage (SL) that overlapped the three ecological niches (human, animal, and marine) was represented at least once in order to maximise diversity within the resulting dataset (10).

We reviewed the draft assemblies and raw reads of the remaining 2896/3255 isolates to select an additional 192 isolates with the aim of maximizing plasmid diversity and capturing clinically relevant features. Raw reads were assessed using SRST2 v0.2.0 (13) with the PlasmidFinder plasmidfinder-db v2023-01-18 (14) and MOB-typer v3.1.5 (15) databases, and draft assemblies were assessed using Abricate v1.0.1 (https://github.com/tseemann/abricate) with the same databases. Isolates were prioritized if they carried a diverse range of replicon markers and/or MOB types. To ensure comprehensive plasmid representation, at least one isolate was selected to represent each replicon and relaxase type, with additional selections for markers appearing across multiple plasmids. Any isolate which contained the only instance of a given plasmid marker in either the raw reads or draft assembly was automatically selected for long-read sequencing. Niche and phylogenetic diversity were also considered in the selection process. Isolates were chosen to represent all three ecological niches, with additional emphasis on traits spanning multiple niches when possible. Representation of multiple KpSC members was also incorporated to ensure broad genomic representation across the dataset. We included isolates where plasmid markers were present in raw reads but absent from draft assemblies, ensuring these sequences were not lost due to assembly artifacts. This resulted in a total of 578 isolates with 1431 plasmids.

### Heavy metal and/or thermoresistance

To identify heavy metal resistance (HMR) genes, we searched the annotated plasmid assemblies for genes listed in the Antibacterial Biocide and Metal Resistance Genes (BacMet) database (http://bacmet.biomedicine.gu.se/index.html) (16). Based on literature searches, we used the following operons to determine presence or absence of heavy metal resistance: Arsenic resistance was determined by the presence of *arsAD*, *arsBCR*, or *arsABCDR* (17–19). Resistance to chromium was defined by the presence of *chrA* or *chrAB* (17,20,21). Mercury resistance was determined by the presence of at least *merAPR* and either *merC*, *merF*, or *merT* (17,22,23). Copper resistance was defined as *pcoABCDRS* (17,24). The *ncrABC* operon was used to define nickel resistance (25). Silver resistance was defined as the presence of at minimum *silABCERS* (17). The *terBCDE* operon was used to determine tellurite resistance (26). The presence of *rcnAR* defined resistance to cobalt and nickel (27). The *zitB* gene indicated resistance to zinc (28). The genes *clpK* and/or *hsp20* were used to determine thermoresistance (29,30).

### Plasmid clustering with Pling

The closed plasmids were clustered with Pling (31), which operates in three main stages: 1) construction of a containment network based on the proportion of sequence shared between two plasmids (i.e. their containment distance), 2) calculation of structural distances using the double-cut and join indel (DCJ-indel) model, and 3) refinement of the plasmid network by identifying hub plasmids.

To identify plasmid pairs that share sufficient sequence for meaningful structural comparison, Pling calculates a *containment distance* between each pair, which is defined as the proportion of the smaller plasmid not aligned to the larger of the two. By default, plasmids with a containment distance ≤0.5 are assigned to the same *community* and retained for further comparison.

Plasmid pairs within the containment threshold are then *integerised*, wherein each sequence is converted to an ordered list of integers representing synteny blocks; shared blocks are assigned the same integer across plasmids. By default, unaligned regions ≥200 bp are treated as *indels* and receive unique identifiers.

Using the integerised representations, Pling computes the *DCJ-indel* distance between plasmids, which reflects the minimal number of structural operations (i.e. rearrangements, including inversions, translocations, insertions, and deletions) needed to convert one plasmid structure into another. By default, plasmids with ≤4 rearrangements are assigned to the same *subcommunity*, which represents the currently evolving unit of those plasmids. These subcommunities were referred to as plasmid clusters.

*Hub plasmids* are defined as highly connected nodes that bridge otherwise unconnected groups. By default, these are identified as plasmids with node degree >10 and the neighbouring node's edge density <0.2, where edge density is the proportion of observed edges among neighbours relative to the maximum possible. Pling iterates over the DCJ-indel network, first removing hub plasmids in the original network, then identifying and removing hubs induced by removal of the first set, etc., until no more hubs remain in the network. Hubs often represent plasmids dominated by transposable elements (TEs), which can inappropriately inflate network connectivity.

5. Håkonsholm F, Hetland MAK, Svanevik CS, Lunestad BT, Löhr IH, Marathe NP. Insights into the genetic diversity, antibiotic resistance and pathogenic potential of *Klebsiella pneumoniae* from the Norwegian marine environment using whole-genome analysis.

International Journal of Hygiene and Environmental Health. 2022 May 1;242:113967.

31. Frolova D, Lima L, Roberts LW, Bohnenkämper L, Wittler R, Stoye J, et al. Applying rearrangement distances to enable plasmid epidemiology with pling. Microbial

Genomics. 2024;10(10):001300.
