## Supplementary figures for "A One Health study of *Klebsiella pneumoniae* species complex plasmids shows a highly diverse and ecologically adaptable plasmidome"

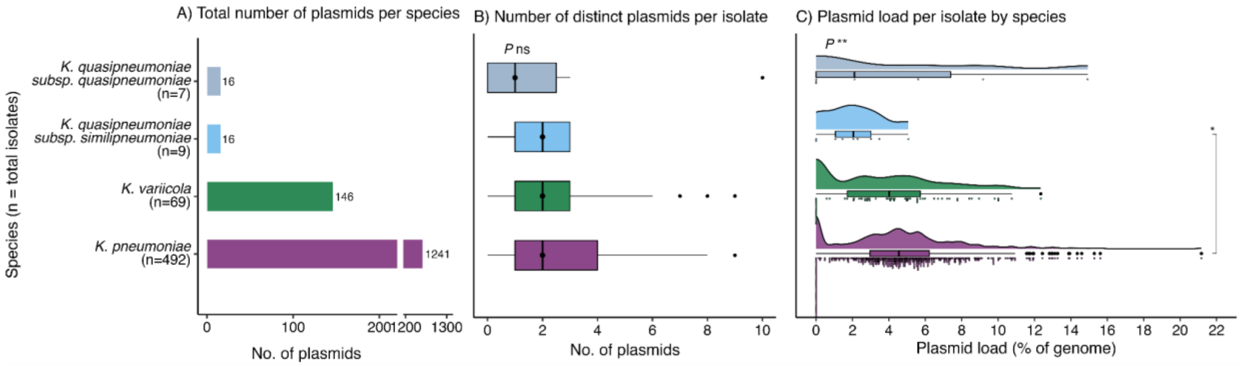


***Figure S1.*** **Plasmid distribution by species. A)** Total number of plasmids per species. **B)** Distribution of the number of distinct plasmid sequences per isolate in each KpSC species, excluding *Klebsiella quasivariicola* (n=1); the black vertical line indicates median value. **C)** Distribution of estimated plasmid load per isolate within each species, excluding *K. quasivariicola*. Statistical comparisons were performed using Kruskal-Wallis (overall) and Mann-Whitney (pairwise) tests. Significance is denoted as follows: *$P<0.05$, ns $P\geq0.05$.

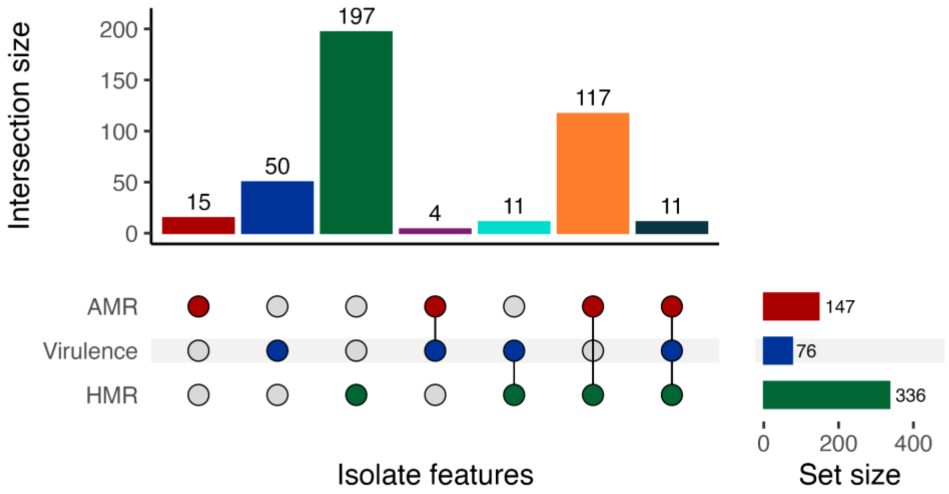


***Figure S2.*** **Overlap of clinically relevant features across whole genomes.** Upset plot showing the overlap of clinically relevant genetic features across 407/578 whole genomes (chromosomes and plasmids). Antimicrobial resistance (AMR) in red, virulence factors in blue, and heavy metal resistance (HMR) in green.

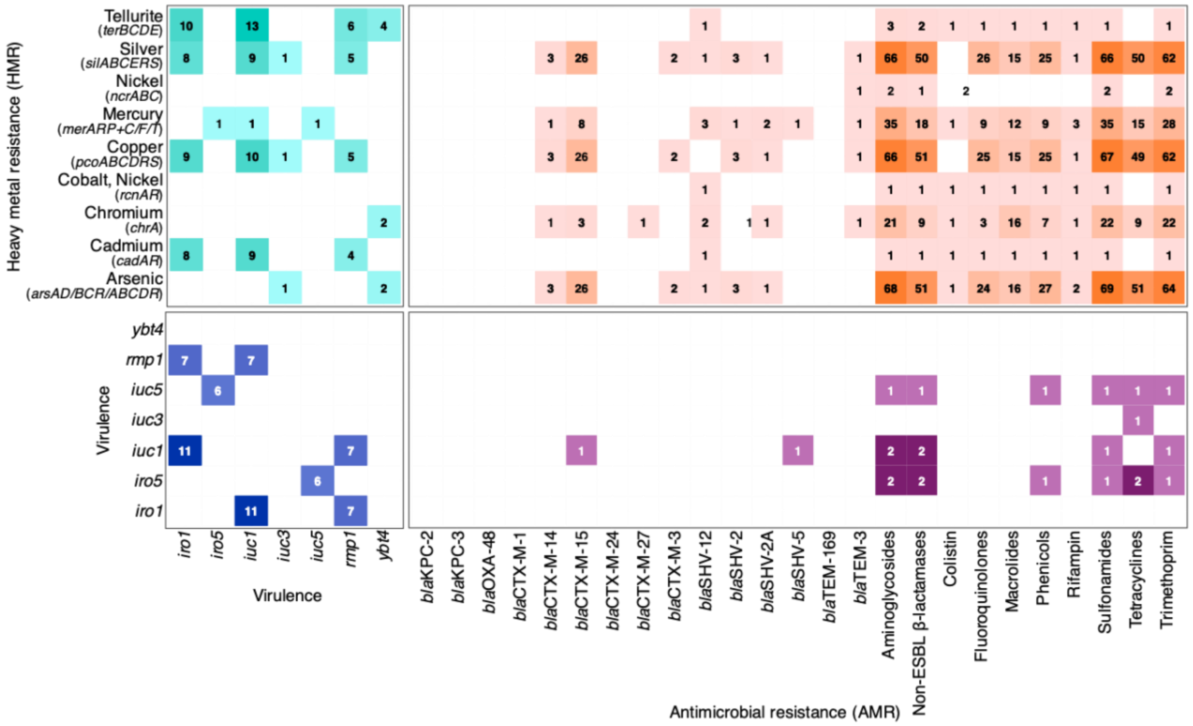


***Figure S3.*** **Co-occurrence of clinically relevant features by plasmid.** Frequency of clinically relevant traits residing on the same plasmid: heavy metal resistance (HMR) and virulence (green), HMR and AMR class or AMR genes (for ESBLs and carbapenemases) (orange), virulence and AMR (purple), and overlapping virulence factors (blue). Not shown: virulence factors or HMR operons that did not overlap with either of the other two categories.

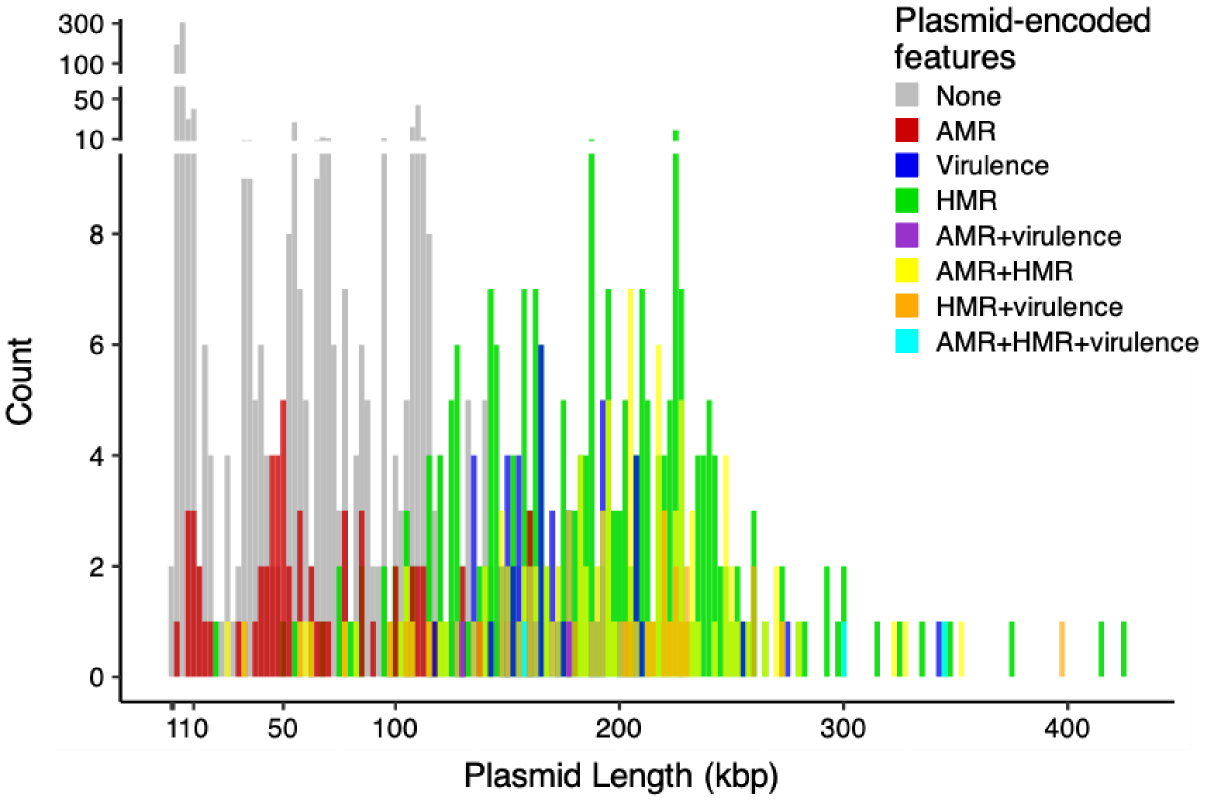


***Figure S4.*** **Distribution of plasmid-encoded features across plasmid lengths**. Each bar shows the number of plasmids in a given length range (x-axis) carrying genes for AMR (red), virulence (blue), AMR and virulence (purple), heavy metal resistance (HMR) and/or thermoresistance (green), AMR and HMR/thermoresistance (yellow), virulence and HMR/thermoresistance (orange), or all three (light blue).

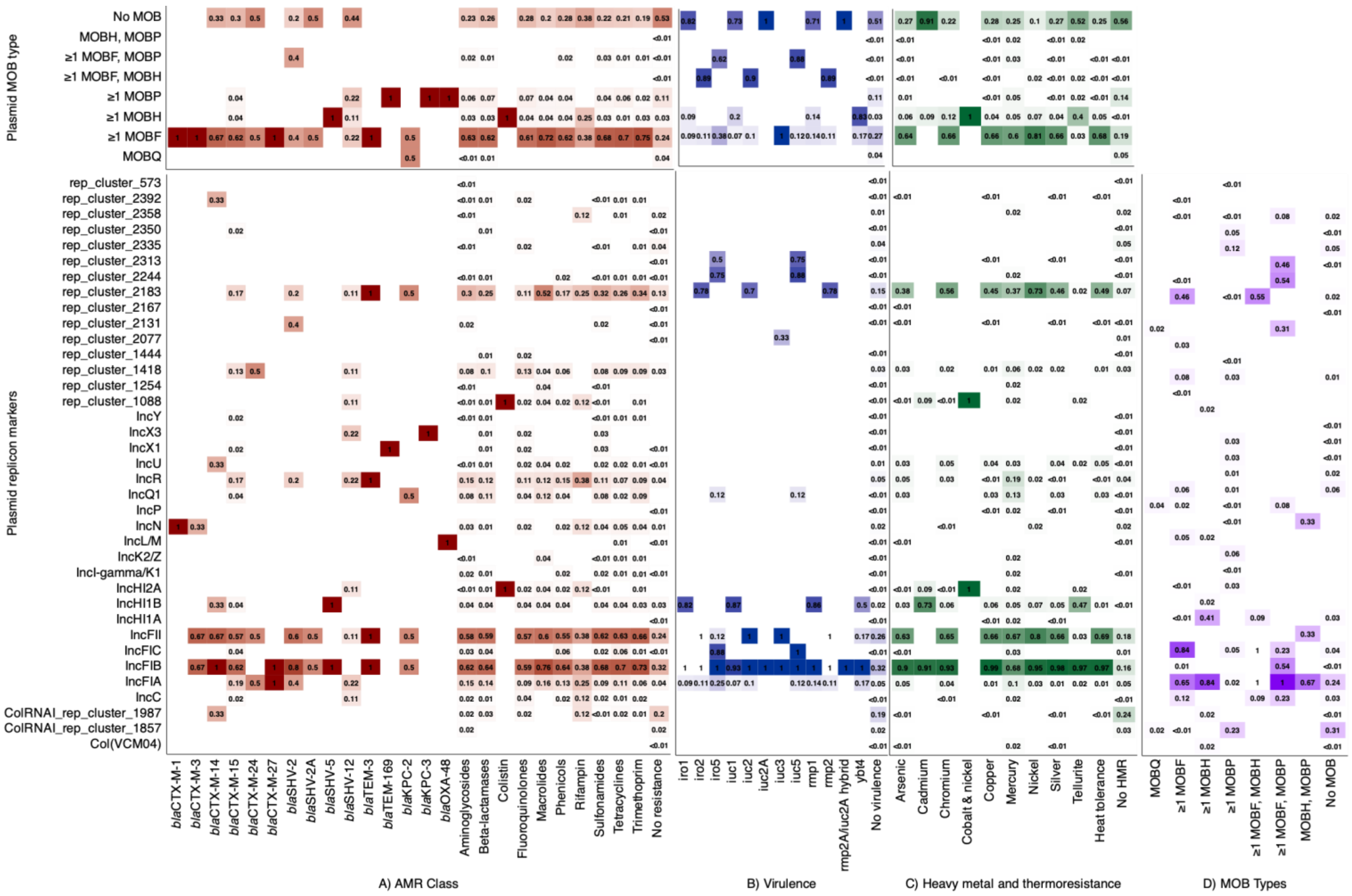


***Figure S5.*** **Overlap of clinically relevant features with replicon and MOB types.** Heatmap with proportion of **A)** antimicrobial resistance (AMR) genes (ESBLs and carbapenemases) or classes, **B)** virulence factors, **C)** heavy metal resistance (HMR) operons, or **D)** MOB types residing on plasmids with plasmid MOB types (top) and replicons (bottom). Not shown: plasmid MOB types and replicons that did not overlap with at least one AMR, virulence, or HMR category.

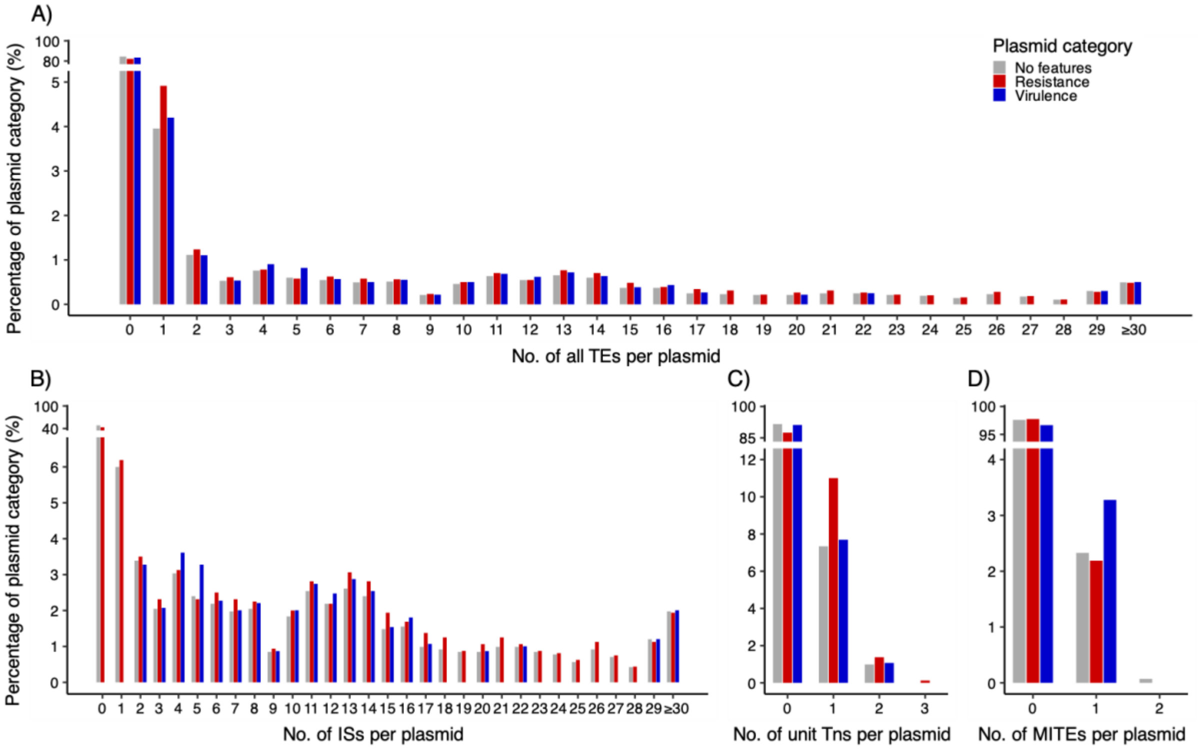


***Figure S6.*** **Prevalence of transposable elements (TEs) across plasmid categories.** Prevalence of **A)** all TE types, **B)** insertion sequences (ISs), **C)** unit transposons (Unit Tns), and **D)** miniature inverted-repeat transposable elements (MITEs) on antimicrobial resistance (AMR) (red), virulence (blue), and non-AMR/virulence (grey) plasmids.

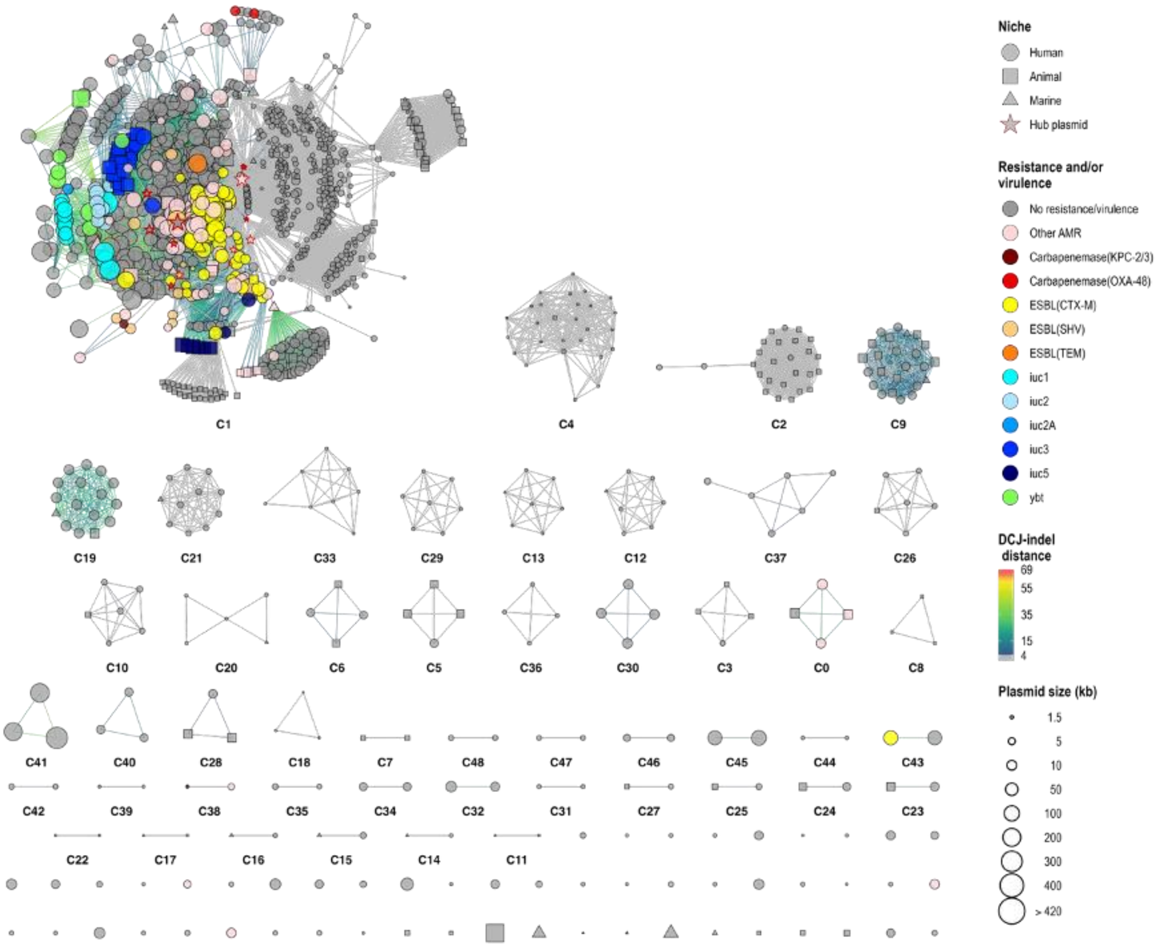


***Figure S7.*** **Full Pling network graph (n=1415 plasmids) including all communities, clusters, singletons, and hub plasmids.** Singletons are not labeled. Hub plasmids (n=13) were only present in the largest community (C1) and are shown as stars with red outlines. Of the 13 hub plasmids, 10 were found in the human niche - two of which encoded ESBLs - and three hub plasmids were found in the marine niche, they did not encode any antimicrobial-, heavy metal- or thermoresistance or virulence.

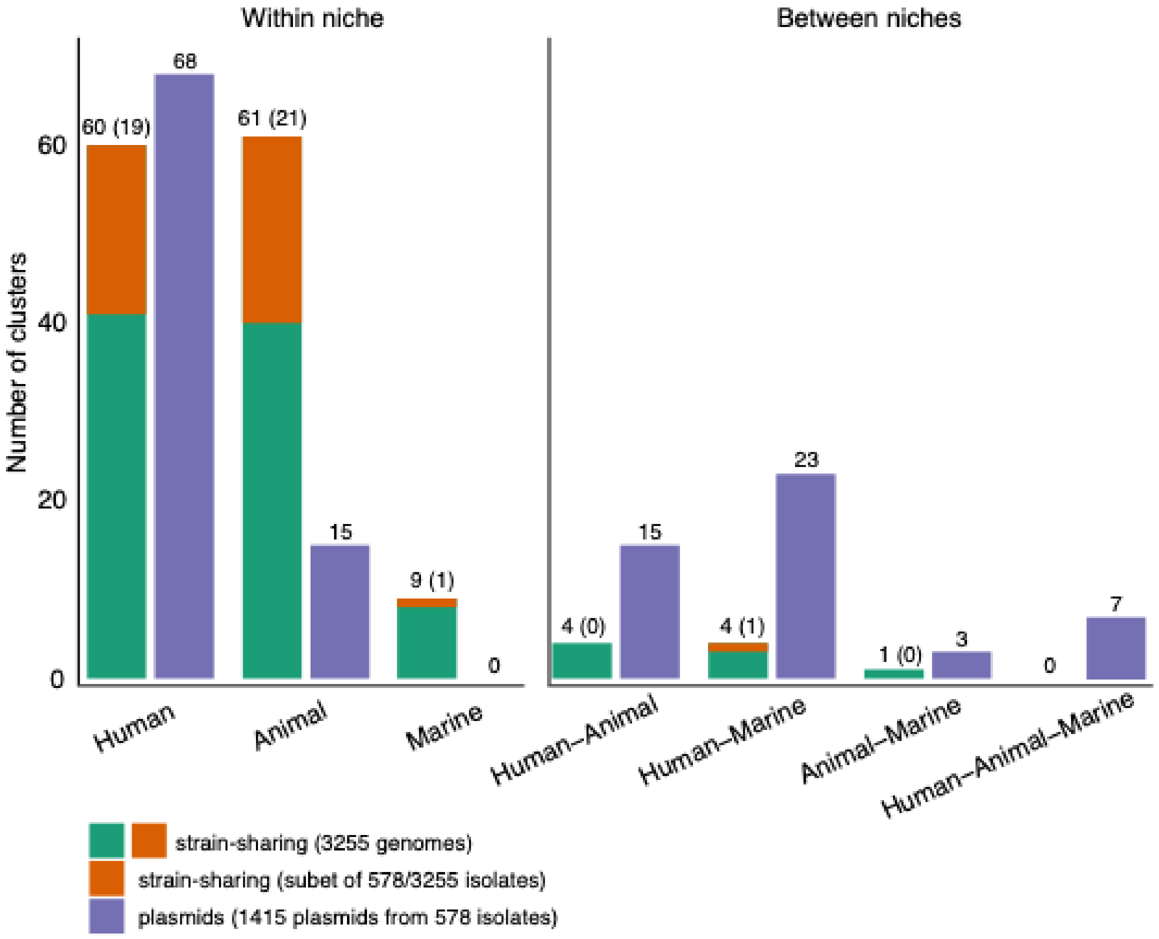


***Figure S8.*** **Cluster strain- versus plasmid- sharing.** Number of Pling clusters that exhibited strain-sharing for all isolates (n=3255) (green) or subset of hybrid-assembled isolates (n=578) (orange), and plasmid sharing (purple) both within each niche (left) and across niches (right).

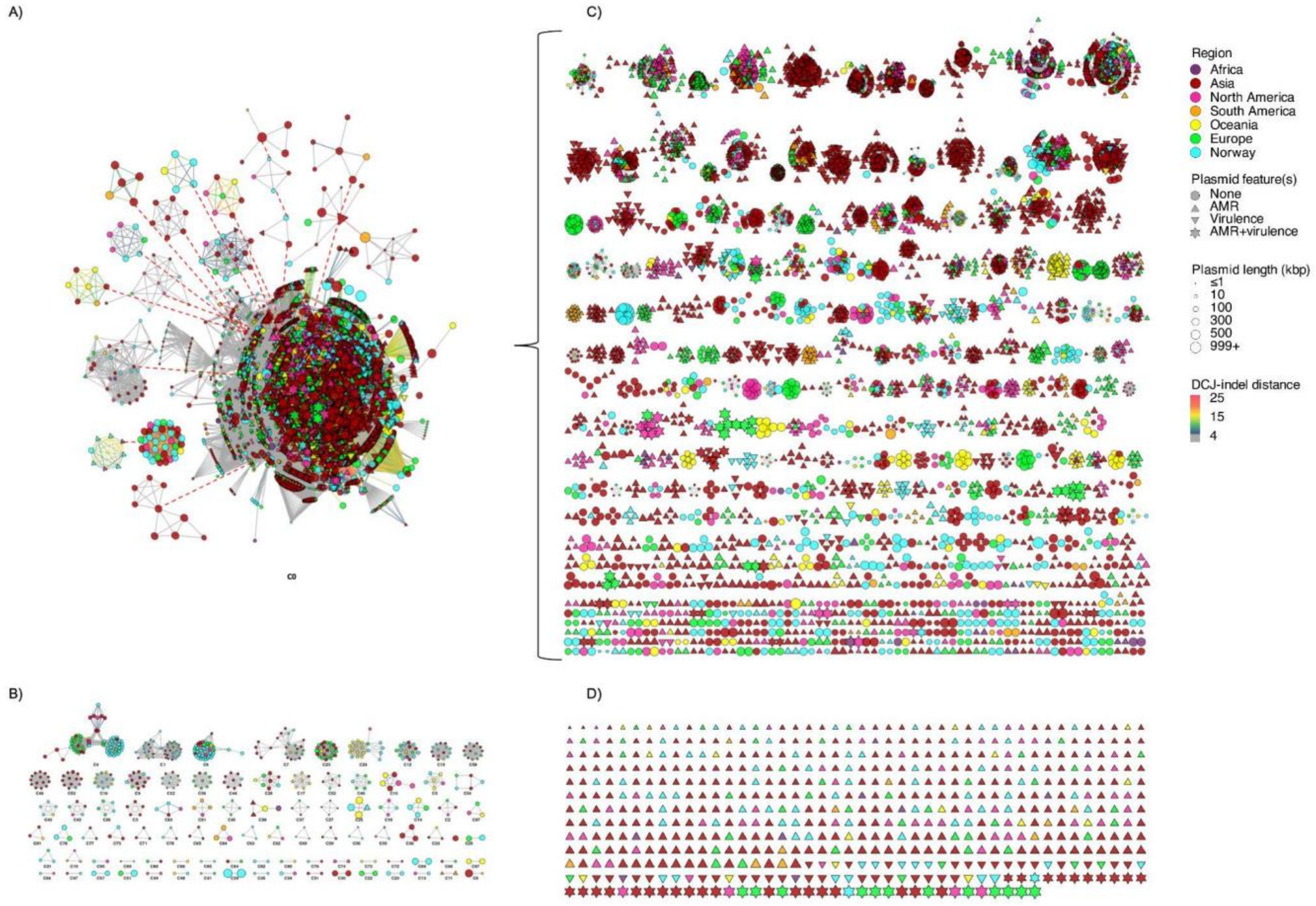


***Figure S9.*** **Pling network graph of global and local plasmids**. **A)** and **B)** Pling plot of communities within the global dataset, excluding 363 hub plasmids (n=9476 plasmids). Red dotted lines indicate connections that would have been made via hub plasmids. **C)** Subview of clusters within the largest Pling community (A) (n=7584 plasmids). **D)** Singletons harbouring antimicrobial resistance (AMR) genes and/or virulence factors (n=564/1544 singletons). Node shape indicates presence/absence of clinically relevant features, node size indicates plasmid length in kbp, node color represents world region, as per inset legend. Edge color indicates DCJ-indel distance between plasmids within the same community, where grey indicates ≤4 rearrangements between plasmids (i.e. plasmids within the same cluster). Not shown: singletons not harbouring AMR genes or virulence factors (n=980/1544 singletons), hub plasmids (n=363; n=224 encoded AMR, n=3 encoded virulence, n=31 encoded AMR and virulence, n=105 did not encode AMR or virulence factors).

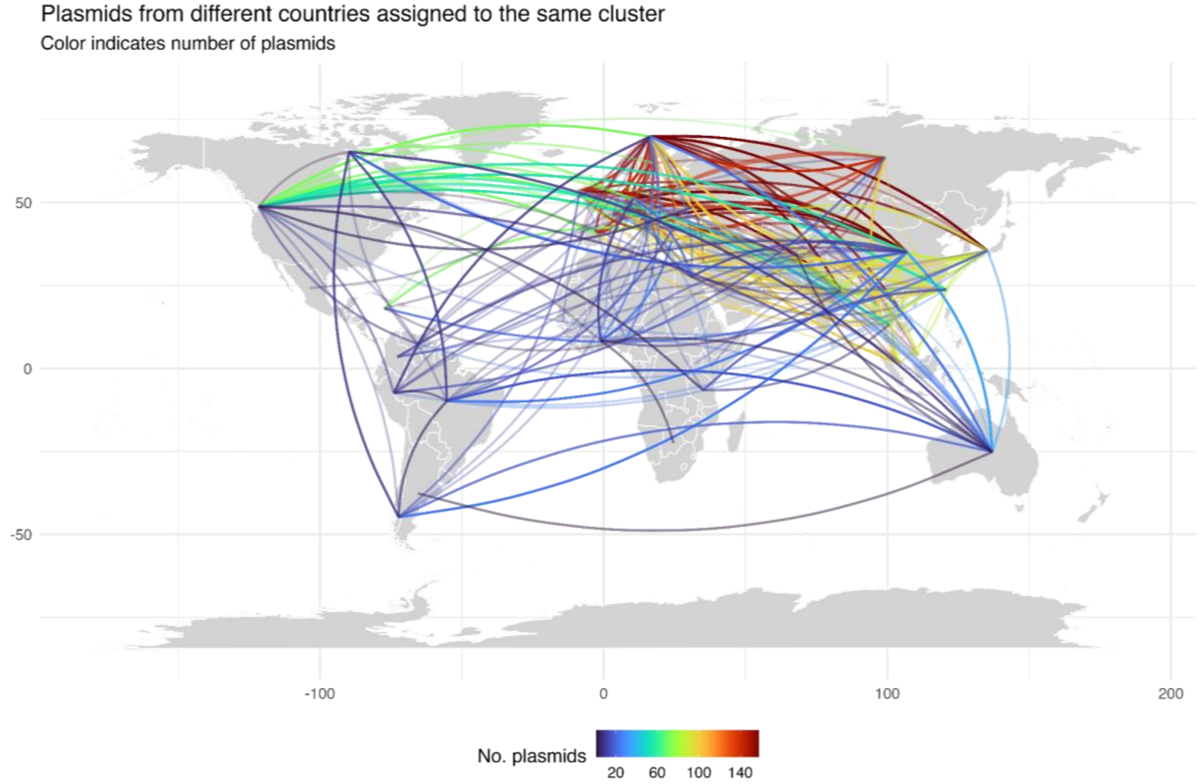


***Figure S10.* Map showing plasmid clusters (n=325) shared across 65 countries and their geographic distribution.** The colour scale indicates the number of plasmids shared between two locations.
